# The pro-apoptotic effect of chronic contractile activity-induced extracellular vesicles on Lewis Lung Carcinoma cells

**DOI:** 10.1101/2024.08.29.610232

**Authors:** Patience O. Obi, Tamiris F.G. Souza, Kirk J. McManus, Adrian R. West, Joseph W. Gordon, Ayesha Saleem

## Abstract

Regular exercise reduces tumor growth *in vivo* and *in vitro*, but the exact mechanisms have yet to be fully elucidated. We have previously shown that chronic contractile activity (CCA) increases the concentration of skeletal muscle-derived EVs, and these in turn increased mitochondrial biogenesis in myoblasts. Here, we hypothesized that skeletal muscle-EVs derived post-CCA will mediate the anti-tumorigenic effects associated with chronic exercise. C2C12 myoblasts were differentiated into myotubes, electrically paced, and EVs isolated from conditioned media from control and CCA myotubes using differential ultracentrifugation. Lewis lung carcinoma (LLC) cells were treated with the total number of control-EVs or CCA-EVs isolated after each day of contractile activity for 4 days. Permeabilized CCA-EVs with or without proteinase K before co-culture with LLC cells were used as controls. Effect of EV treatment on cell count, viability, apoptosis, senescence, migration, and mitochondrial content was measured. CCA-EV treatment reduced cell count by 18% and cell viability by 6% *vs.* control-EVs. CCA-EVs increased the incidence of apoptotic hallmarks: DNA fragmentation by 13%, Annexin V+/PI+ cells by 21%, and the expression of pro-apoptotic Bax (by 25%) and Bax/Bcl-2 ratio (by 60%) *vs.* control-EVs. CCA-EVs increased number of senescent cells by 29%, and senescence markers, HMGB1 (by 49%) and p16 (by 92%) *vs.* control-EVs. When CCA-EVs were pretreated with Triton X-100 with or without proteinase-K, the increase in apoptosis and senescence was abrogated, confirming the effect is due to intact EVs and likely through EV membrane-proteins. CCA-EVs did not have any effect on cell migration and mitochondrial content *vs.* control-EVs. This study illustrates for the first time the potential of CCA-induced skeletal muscle-EVs in mediating anti-tumorigenic effects traditionally linked with chronic exercise.

## Introduction

Lung cancer is the leading cause of cancer-related deaths worldwide, accounting for about 1 in 5 of all cancer deaths among men and women each year [1]. There are two main types of lung cancer: non-small cell lung cancer (NSCLC) and small cell lung cancer (SCLC). NSCLC accounts for about 80-85% of lung cancer cases, and this type of lung cancer usually grows at a slower rate than SCLC. NSCLC is further sub-divided into three main types: adenocarcinoma, squamous cell carcinoma, and large cell carcinoma [2]. Standard treatment options include chemotherapy, radiotherapy, surgery, immunotherapy, targeted therapy, and palliative procedures [3], [4], but there is no cure for lung cancers to date. Adjuvant therapies including exercise are often administered after the main treatment to help manage cancer [5]–[7].

Exercise is associated with reduced risk of several types of cancers including lung cancer [8], [9]. Exercise not only lowers the risk of cancer, but improves the quality of life of cancer patients by increasing the efficacy of main or adjuvant cancer treatments [5], [7]. Furthermore, pre-clinical *in vivo* and *in vitro* studies have reported that endurance exercise can inhibit tumor incidence, growth and metastasis [10], [11]. Possible mechanisms underlying this beneficial effects of exercise include those involving immune surveillance (natural killer cells, macrophages and T-regs) [10], [12] and myokines such as irisin, muscle-derived oncostatin (OSM) and secreted protein acidic and rich in cysteine (SPARC) have been implicated, but the exact mechanism(s) by which exercise exerts anti-cancer effects have yet to be fully elucidated [11], [13], [14]. Strong evidence suggests that the systemic adaptative effects of exercise are potentiated in part by the release of factors known as myokines during exercise from skeletal muscle [15]. Myokines are usually proteins that can travel to different organs in the body and act in an autocrine, paracrine, or endocrine manner. Examples include interleukin (IL)-6, IL-10, IL-15, irisin, meteorin, myostatin, among others [15]. Myokines can be released from the skeletal muscle into the circulation by themselves or they can be packaged within extracellular vesicles (EVs) [15], [16].

EVs are small lipid bilayer-delimited particles that are produced from all living prokaryotic and eukaryotic cells, and play a central role in cellular communication. EVs have been traditionally divided into: i) exosomes (30 – 150 nm in diameter), formed by the exocytosis of intraluminal vesicles upon fusion of multivesicular bodies (MVBs) with the plasma membrane, ii) microvesicles (100 – 1000 nm), that are formed by the outward budding of the plasma membrane, and iii) apoptotic bodies (500 – 5000 nm) which are formed from the outward blebbing of an apoptotic cell membrane [17], [18]. More recently, the Minimal Information for Studies of EVs (MISEV) 2023 guidelines recommend that EVs be classified based on size, (where small EVs or sEVs are <200 nm and large EVs are >200 nm), biochemical composition, and cell of origin [18]. EVs contain biochemical cargo, including DNA, mRNA, miRNA, proteins, lipids, and metabolites and can transfer them from parent to recipient cells [16]. There is growing interest in whether EVs play a role in mediating the beneficial systemic effects of exercise. Several studies have shown that exercise increases systemic EV concentration, and modifies cargo components [19]–[22], but substantial evidence investigating the biological activity of exercise-derived EVs in cancer is lacking. A recent study showed that plasma-EVs isolated from exercised rats delayed the progression of prostate cancer [23], but the exact mechanism underlying this effect is unknown. Moreover, this work was done using EVs isolated from plasma (circulation), which comprise a heterogenous mixture of vesicles derived from platelets, erythrocytes, endothelial cells, leukocytes, [22], [24], and comparatively small proportion from skeletal muscle [25], [26]. Given the central role of the skeletal muscle in releasing myokines that mediate the effects of exercise, we hypothesized that skeletal muscle-derived EVs (Skm-EVs) isolated post-exercise will have an anti-tumorigenic effect on lung cancer cells. We have recently demonstrated that chronic contractile activity (CCA) increases the concentration of Skm-EVs, and these in turn increased mitochondrial biogenesis in C2C12 myoblasts [27]. Since Skm-EVs isolated post-CCA had beneficial effects on healthy cells, we believe the same EVs would have some comparable effects on non-healthy cells, in this case, lung cancer cells.

In this study, we investigated the biological activity of Skm-EVs derived post-exercise on NSCLC cells CCA to mimic chronic endurance exercise *in vitro*. Utilizing this well-characterized model of exercise, we assessed the effect of CCA-induced Skm-EVs on Lewis lung carcinoma (LLC) cells, a murine NSCLC cell line. We hypothesized that Skm-EVs isolated after CCA would suppress cell growth by inducing apoptosis and/or senescence in LLC cells. To evaluate this hypothesis, we purified EVs from conditioned media of both non-contracted control myotubes and from myotubes post-CCA. Both control and CCA-derived EVs were co-cultured with LLC cells, and the subsequent effect on cell growth, viability, apoptosis, and senescence in recipient cells was measured.

## Methods

### Cell culture

Murine C2C12 myoblasts (500,000 cells/well) were seeded in a six well plate pre-coated with 0.2% gelatin, and grown in fresh Dulbecco’s Modification of Eagle’s Medium (DMEM; Sigma-Aldrich) supplemented with 10% fetal bovine serum (FBS; Gibco/ThermoFisher Scientific) and 1% penicillin/streptomycin (P/S) (growth media). Cells were grown at 37 °C in 5% CO_2_ incubator for 24 h. When myoblasts reached approximately 90-95% confluency, the growth media was switched to differentiation media (DMEM supplemented with 5% heat-inactivated horse serum (HI-HS; Gibco/ThermoFisher Scientific) and 1% P/S) for 5 days to differentiate myoblasts into myotubes. Lewis Lung Carcinoma (LLC) cells, as a model of NSCLC cells, were obtained from the American Type Culture Collection (ATCC; CRL-1642) and maintained in media containing DMEM supplemented with 10% FBS, and 1% P/S. All cells were grown at 37 °C in 5% CO_2_ incubator.

### Chronic contractile activity-derived EVs and EV co-culture with LLC cells

After five days of differentiation, myotubes were divided into control (CON) and chronic contractile activity (CCA) groups. CCA was performed using electrical pulse stimulation as previously described [27]. Briefly, the CCA plate was stimulated using the C-Pace EM Cell culture stimulator, with C-dish and carbon electrodes (IonOptix, Milton, MA, United States) while the CON plate had the C-dish placed on top but without the carbon electrodes. Myotubes were subjected to CCA: 1 Hz (2 ms pulses), at 14 V for 3 h/day for four consecutive days. After the first bout of contractile activity, spent media was exchanged with exosome-depleted differentiation media (DMEM supplemented with 5% exosome-depleted HI-HS and 1% P/S) for both CON and CCA groups and myotubes left to recover for 21 h. After recovery, 10-12 mL of conditioned media was collected from the cells and used for EV isolation. This process was repeated after each bout of contractile activity for 4 days. The EVs collected after the first bout of contractile activity (labelled as Day 1 EVs) was then used to treat LLC cells, and the same was done for EVs collected after each bout of contractile activity. LLC cells were seeded at a density of 90,000 cells/well in 6-well plates, or at 5000 cells/well in 24 well plates in growth media and allowed to adhere for 4 h. Subsequently, cells were washed with PBS and exosome-depleted media (DMEM supplemented with 5% exosome-depleted HI-HS and 1% P/S) was added to cells before treatment with EVs. The EV treatment was performed after each day of contractile activity, (i.e. Day 1 EVs were used to treat the cells on the first day of treatment), and this was done continuously for four days. Exosome-depleted HI-HS was used to remove any confounding effects of EVs found naturally in horse serum. To deplete horse serum of exosomes/sEVs, we diluted it to 10% with DMEM, spun at 100,000*xg* for 18 h and filtered with 0.22 µm filter before making the media, which we labelled as exosome-depleted differentiation media.

### Isolation of EVs by differential centrifugation

EV isolation by differential ultracentrifugation (dUC) was performed according to the protocol by Théry et al. [28] and as previously described [27]. Briefly, conditioned media (CM) from each six well plate (12 mL) was collected and centrifuged at 300*xg* for 10 min at 4 °C to pellet dead cells (Sorvall™ RC 6 Plus Centrifuge, F13-14 fixed angle rotor), followed by centrifugation at 2000*xg* for 10 min at 4 °C to remove cell debris. The resulting supernatant was centrifuged at 10,000*xg* for 30 min at 4 °C to remove large vesicles. Using an ultracentrifuge (Sorvall™ MTX 150 Micro-Ultracentrifuge, S58-A fixed angle rotor), the supernatant was centrifuged at 100,000*xg* for 70 min at 4 °C to obtain the exosome/sEV pellet, after which the resulting supernatant, labelled as EV-depleted conditioned media (EV-dep) was collected and stored for later use. The exosome/sEV pellet was resuspended in 1 mL PBS and centrifuged again at 100,000*xg* for 70 min at 4 °C. After centrifugation, the final exosome/sEV pellet was resuspended in 50 µL PBS and used for subsequent analysis.

### Trypan blue exclusion assay

LLC were treated with CON-EVs and CCA-EVs as described above. Cell proliferation and viability was measured with trypan blue exclusion as we described previously [27]. After the last day of EV treatment, LLC cells were washed with PBS, trypsinized and centrifuged at 1000*xg* for 5 min. Cells were resuspended in 1 mL of growth media, stained with a 1:25 dilution of Trypan blue (Sigma-Aldrich, T8154) and counted with a hemocytometer (Hausser Scientific Bright-Line Hemacytometer, Sigma-Aldrich). Total number of cells was counted and expressed per mL of growth media. The number of viable cells were counted and the cell viability was obtained by dividing the number of viable cells by the total number of cells.

### MTT assay

Cell viability was also measured by Thiazolyl Blue Tetrazolium Bromide (MTT; Sigma-Aldrich, M5655) assay as previously [27]. LLC cells were seeded at 5000 cells/well in 96-well plates with exosome-depleted media in triplicates, and treated with CON-EVs or CCA-EVs for four days after each bout of contractile activity. Since we used EVs isolated from 12 mL of CM to treat 90,000 cells/well in previous assays, we divided the EVs into 18 equal portions to treat 5000 cells/well. After treatment, 20 µL of MTT Reagent 5 mg/mL was added and incubated for 3 h. The yellow tetrazolium MTT was reduced to formazan crystals by intracellular NAD(P)H-oxidoreductases. 150 µL of DMSO was added and mixed gently to solubilize the formazan crystals. The assay was quantified by spectrophotometry at the absorbance of 540 nm using a microplate spectrophotometer (BioTek, Epoch). Cell viability was calculated as a ratio of absorbance of sample to the absorbance of untreated control.

### Preparation of cell enzyme extracts

LLC cells were harvested after the last day of treatment using a cell scraper (Corning®), resuspended in 50 µL of enzyme extraction buffer (100 mM Na-K-phosphate, 2 mM EDTA, pH 7.2), sonicated 3 x 3 seconds (s) on ice, subjected to repeated freeze-thaw cycles, vortexed vigorously for 5 s and centrifuged at 14,000 rpm as before [29]. The supernatant was collected and used to measure protein concentration and yield, cytochrome *c* oxidase (COX) enzyme activity and Western blotting.

### Protein concentration and yield

Protein concentrations of whole cell lysates were determined using Pierce™ BCA protein assay kit (ThermoFisher Scientific) following the manufacturer’s instructions as described before [30]. Briefly, a serial dilution of 2 mg/mL bovine serum albumin (BSA) in sterile water was prepared and used as standard. 25 µL of each standard or sample was added in duplicates into a 96-well microplate, followed by the addition of 200 µL working reagent to each well. Samples were incubated at 37 °C for 30 min and the absorbance was measured at 562 nm using a microplate spectrophotometer (BioTek, Epoch). Protein yield was determined by multiplying the protein concentration with the volume of cell lysate.

### MitoTracker Red staining

LLC were treated with CON-EVs and CCA-EVs as described above. After the last day of treatment, LLC cells were washed with PBS, trypsinized and 10,000 cells/well were re-seeded in a 24-well plate. Post-adherence for 24 h, treated LLC cells were incubated with MitoTracker Red CMXRos (Cell Signaling Technology, 9082) and Hoechst 33258 (Sigma-Aldrich, 94403) at final concentrations of 50 nM and 3 µg/mL, respectively, under normal culture conditions for 30 min. Following incubation, cells were washed with PBS, and medium was switched to growth media. Cells were visualized using the Olympus IX70 inverted microscope (Toronto, ON, Canada) with NIS Elements AR 3.0 software (Nikon Instruments Inc., Melville, NY, USA). Three representative images per well were taken, and the fluorescence intensity was quantified using ImageJ software (NIH, Bethesda, MD, USA), where the free hand tool was used to draw the region of interest (ROI) from five LLC cells for each representative image, and mean intensity for each was averaged to get the total fluorescence intensity.

### COX activity

We measured mitochondrial COX activity, a gold standard marker of mitochondrial biogenesis, in the LLC enzyme extracts as previously described [31]. Briefly, enzyme extracts were added to a test solution containing fully reduced cytochrome *c* (Sigma-Aldrich). COX activity was determined as the maximal rate of oxidation of fully reduced cytochrome *c* measured by the change in absorbance over time at 550 nm using an imaging multi-mode reader (BioTek, Cytation 5) at 30 °C. Protein concentration was determined using the Pierce™ BCA protein assay kit as described above, and COX activity was normalized by total protein content.

### Western blotting

10 µg of total protein from whole cell extracts were resolved on a 12% SDS-PAGE gel and subsequently transferred to nitrocellulose membranes. Membranes were blocked for 1 h with 5% skim milk in 1× Tris-buffered saline-Tween 20 solution (TBST) at room temperature, followed by incubation with primary antibodies in 1% skim milk overnight at 4 °C. The following primary antibodies were used: mouse anti-p53 (2524, Cell Signaling, 1:500), rabbit anti-Apoptosis Inducing Factor (AIF) (5318, Cell Signaling, 1:1000), rabbit anti-caspase-3 (9662, Cell Signaling, 1:1000), rabbit anti-cleaved caspase-3 (9661, Cell Signaling, 1:1000), rabbit anti-Bcl-2 (3498, Cell Signaling, 1:1000), rabbit anti-Bax (2772, Cell Signaling, 1:1000), rabbit anti-cytochrome *c* (AHP2302, Bio-Rad Laboratories, 1:250), rabbit anti-High mobility group box 1 (HMGB1) (6893, Cell Signaling, 1:2000), mouse anti-p16 (SAB4500072, Sigma-Aldrich, 1:500), rabbit anti-cyclin D1 (2922, Cell Signaling, 1:1000), and mouse OXPHOS cocktail (45-8099, Invitrogen, 1:500). Subsequently, membranes were washed three times for 5 min with TBST, followed by incubation with anti-mouse (A16017, ThermoFisher) or anti-rabbit (A16035, ThermoFisher) IgG horseradish peroxidase secondary antibody (1:1,000-10,000) in 1% skim milk for 1 h at room temperature, after which the membrane is washed again three times for 5 min each with TBST. Membranes were visulaized using enhanced chemiluminescence detection reagent (Bio-Rad Laboratories), and the films were scanned using the ChemiDoc^TM^ MP Imaging System (Bio-Rad Laboratories). The intensity of bands was quantified using the Image Lab Software (Bio-Rad Laboratories) and corrected for loading using ponceau S staining.

### Treatment of EVs with Proteinase K/Triton X-100

To delineate the specificity of effects were attributable to CCA-EV proteomic cargo, CCA-EVs were treated with 0.1% Triton X-100 for 30 min at room temperature, and incubated with or without 10 µg/mL proteinase K (ProK; Sigma-Aldrich, P2308) for 1 h at 37 °C as detailed before [27]. ProK activity was inhibited by adding 5 mM phenylmethylsulfonyl fluoride (PMSF; Sigma-Aldrich, 10837091001) for 10 min at room temperature. The samples were further processed by addition of 6 mL PBS followed by ultracentrifugation at 100,000*xg* for 70 min at 4 °C to eliminate Triton X-100 before co-culture with cells. After centrifugation, the final pellet (not visible) was resuspended in 50 µL PBS, and used to treat LLC cells.

### Annexin V-FITC/Propidium Iodide (PI) staining assay

The number of apoptotic cells were detected with the annexin V-FITC apoptosis detection kit (Sigma-Aldrich, 11858777001). Annexin V-FITC/PI staining shows the number of cells undergoing early and late apoptosis, as well as necrotic and viable cells. LLC cells were seeded at 5000 cells/well in 24-well plates with exosome-depleted media, and treated with CON-EVs or CCA-EVs for four days after each bout of contractile activity. CCA-EVs were also pretreated with Triton X-100 as described above before co-culture with LLC cells. After treatment, cells were washed with PBS, trypsinized and centrifuged at 1000*xg* for 5 min. Cells were resuspended in 100 µL incubation buffer before adding 2 µL of FITC-conjugated Annexin V and 2 µL of PI, and incubated for 10 min at room temperature in the dark, according to the manufacturer’s instructions. All samples were measured for the Annexin V-FITC/PI fluorescence intensity by flow cytometry using the Attune NxT Acoustic Focusing Flow Cytometer. Each independent experiment was performed in triplicates.

### Cell Death ELISA assay

Cell death was detected using the Cell Death Detection ELISA Kit (Roche, 11544675001), which measures the level of DNA fragmentation, a well-documented hallmark of apoptosis [32]. This assay allows the detection of mono- and oligonucleosomes generated from apoptotic cells using antibodies directed against DNA and histones. Briefly, LLC cells were seeded at 5000 cells/well in 24-well plates and co-cultured with EVs as described above. CCA-EVs were also pretreated with Triton X-100 and proteinase K before co-culture with LLC cells. After treatment, cells were washed with PBS, trypsinized and centrifuged at 1500*xg* for 5 min. Cells were resuspended in 500 µL of incubation buffer and incubated for 30 min at room temperature to allow for lysis. Cell lysate were then centrifuged at 20,000*xg* for 10 min, and 400 µL of the supernatant (cytoplasmic fraction) was carefully collected. The resulting supernatant was diluted (1:10) with incubation buffer and used for the ELISA assay according to the manufacturer’s instructions. Absorbance (Ab) was measured at 405 nm and 490 nm using a microplate spectrophotometer (BioTek, Epoch), and the enrichment factor (EF) in small nucleosomes was calculated with the formula EF = Ab(treated)/Ab(PBS). The experiment was repeated independently in triplicates.

### Senescence-Associated β-Galactosidase assay

Senescence-associated β-galactosidase (SA-β-gal) activity is a classic maker of senescence in cellular biology [33]. Senescence was measured using Senescence Cells Histochemical Staining Kit (Sigma-Aldrich, CS0030) according to the manufacturer’s instructions. This assay detects senescent cells based on a histochemical stain for β-galactosidase activity at pH 6. Briefly, LLC cells were seeded at 5000 cells/well in 24-well plates and co-cultured with EVs as described above. CCA-EVs were also pretreated with Triton X-100 and proteinase K before co-culture with LLC cells. After treatment, cells were washed with PBS, trypsinized and 10,000 cells/well were re-seeded in 24-well plates with growth media. After 24 h, the cells were fixed using 20% formaldehyde for 10 min, after which they were washed with PBS, stained with the staining solution to detect SA-β-Gal activity and left overnight in a non-CO_2_ incubator. The next day, representative images were taken at 4X and 10X magnification with 3 images per magnification using fluorescence microscopy (Cytation 5 Cell Imaging Multi-Mode Reader) and quantified using ImageJ. The percentage of SA-β-gal positive cells was evaluated by dividing the number of SA-β-gal positive cells by the total number of cells for each image. Each experiment was performed in triplicates.

### Wound healing assay

The migratory ability of LLC cells was determined by wound healing (scratch) assay as described previously [34]. LLC cells were seeded at 5000 cells/well in 24-well plates and co-cultured with EVs as described above. CCA-EVs were also with pretreated Triton X-100 and proteinase K before co-culture with LLC cells. After treatment, cells were washed with PBS, trypsinized and re-seeded in 24-well plates with growth media. After 24 h, the cells were treated with 10 µg/ml of mitomycin C (Sigma-Aldrich, CM4287) for 1 h at 37 °C to prevent proliferation. A sterile 1000 µL micropipette tip was used to make linear wounds in the middle of the wells. The cells were then washed twice with PBS and fresh growth media was added. To monitor the cell migration, the cells were imaged in the same position of the wound using a microscope (EVOS™ XL Core Imaging System) with 10X objective at 0 h (time of wounding), 6 h and 24 h. The wound closure was evaluated by measuring the difference in the area of the wounds at the different time points using ImageJ. The formula used was wound closure (%) = (initial wound area − wound area of specified time)/ initial wound area × 100%. Each independent experiment was performed in triplicates.

### EV fluorescent labeling

For confirmation of EV uptake in cells, EVs were labelled with MemGlow™ 488 Fluorogenic Membrane Probe (Cytoskeleton) as previously described [27]. EVs were incubated with 200 nM MemGlow labelling solution for 1 h at room temperature keeping it protected from light. To remove unbound dye from the solution, EVs were diluted with filtered PBS and centrifuged at 100,000*xg* for 70 min at 4 °C, after which the final labelled EVs were collected. To perform EV cell uptake experiments, 50 µL of EVs (from 10 mL of conditioned media) were labelled with 200 nM MemGlow 488, and co-cultured with 20,000 LLC cells/well in an 8-well chamber slide for 24 h. Subsequently, cells were fixed with 3.7% paraformaldehyde (PFA), stained with 66 µM of Rhodamine phalloidin and 2-(4-amidinophenyl)-1H-indole-6-carboxamidine (DAPI), and then used for confocal imaging (LSM 700 confocal microscope, Zeiss, St. Louis, USA).

### Statistical analysis

All data were analyzed using unpaired Student’s t-test, one-way ANOVAs, or two-way ANOVAs. Multiple comparisons in the one-way ANOVA were corrected using Holm-Šídák post hoc test, or if the data did not pass normality, with Dunn’s post hoc test. Normality was measured using a combination of tests: D’Agostino-Pearson, Anderson-Darling, Shapiro-Wilk and Kolmogorov-Smirnov when possible. Multiple comparisons in the two-way ANOVA were corrected using Bonferroni’s post hoc test. Individual data points are plotted, with mean ± standard error of mean (SEM) shown as applicable. All graphs were created using GraphPad-Prism software (version 10.1.2, GraphPad, San Diego, CA, USA). Significance was set at p ≤ 0.05. Exact p values for significant or close to statistically significant results is shown. An n = 3-7 was conducted for all experiments.

## Results

### CCA-EVs inhibits non-small cell lung cancer cell viability

The characterization of myotubes and EVs isolated from control and chronically stimulated myotubes has been previously published by our group [27]. The overall study design is shown in **Fig. 1**. The effect of CCA-EVs on cell viability was evaluated by trypan blue exclusion and MTT assays. The results showed that CCA-EVs inhibited the growth of LLC cells by decreasing the total cell number by 18% (p=0.0071, n=6, **Fig. 2A**) as well as the percentage of viable (live) cells compared to CON-EVs by 6% (p=0.0383, n=6, **Fig. 2B**). Cell viability as measured by the MTT assay however remained unchanged (**Fig. 2C**), along with total protein yield from LLC cells in CCA-EVs *vs*. CON-EV treated cells (**Fig. 2D**).

**Figure 1.**
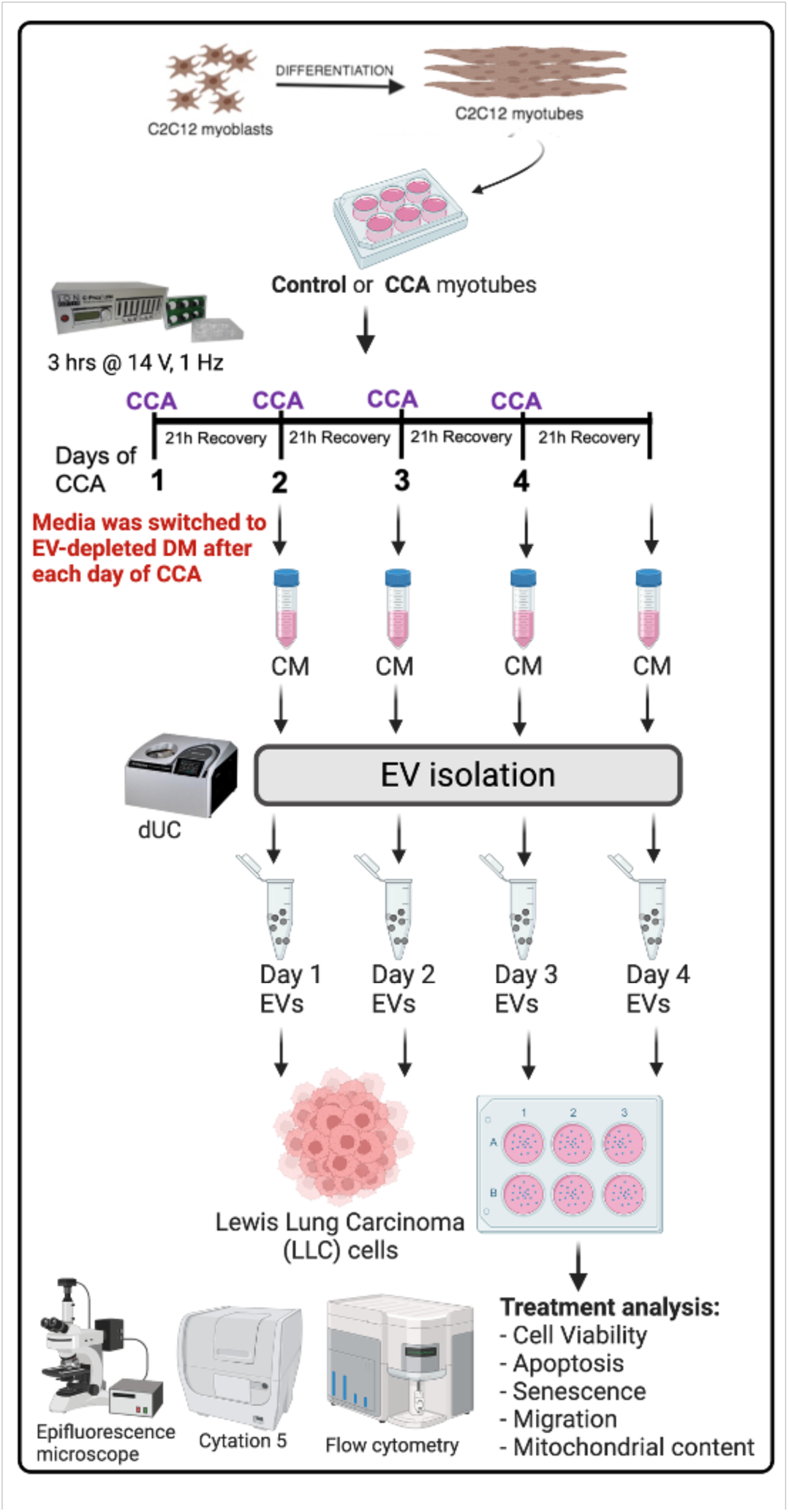
Overall study design. C2C12 myoblasts were fully differentiated into myotubes (MTs). MTs were divided into control (CON) and chronic contractile activity (CCA) plates. CCA-MTs were electrically paced for 3h/day x 4 days at 14 V to mimic chronic endurance exercise *in vitro* using IonOptix ECM. After the first bout of contractile activity, media was replaced with exosome-depleted differentiation media in both CON-MT and CCA-MT, and cells were allowed to recover for 21 h. Conditioned media from control or stimulated myotubes was used to isolate EVs via dUC, and EVs were co-cultured with Lewis Lung Carcinoma (LLC) cells. This process was done after each bout of contractile activity from Day 1-4. After the last day of treatment, cells were collected and assessed for cell viability, apoptosis (cell death), senescence, cell migration and mitochondrial content. Figure created with BioRender.com

**Figure 2.**
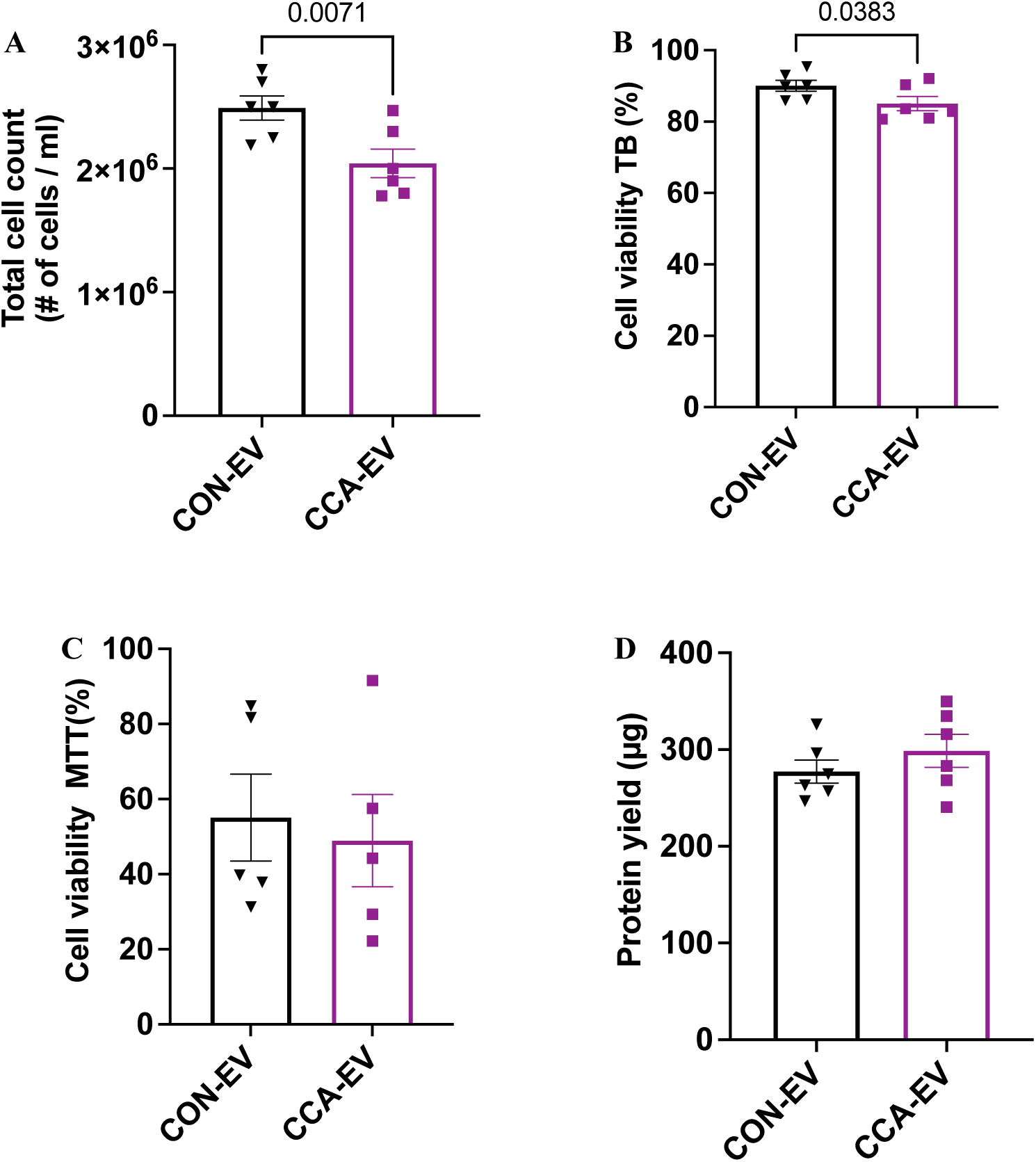
CCA-EVs decreases cell count and viability in LLC cells. EVs were isolated after each bout of contractile activity as previously described, and co-cultured with LLC cells. After the last day of treatment, cells were collected and assessed for; **(A)** Total cell count **(B)** Cell viability using trypan blue (TB), **(C)** Cell viability using MTT, and **(D)** Protein yield in LLC cells treated with CCA-EVs *vs*. CON-EVs. Data were analyzed using an unpaired Student’s t-test and expressed as scatter plots with mean (n=6). Exact p values for significant (p<0.05) results are shown. The p-values for non-significant data are not shown.

### CCA-EVs induces apoptosis in non-small cell lung cancer cells

We examined the effect of CCA-EVs on induction of apoptosis using Annexin V-FITC/PI staining, Cell Death Detection ELISA and expression of apoptotic proteins. CCA-EVs induced cell death in LLC cells by increasing the number of double-stained cells (late-stage apoptotic cells) by 28% compared to cells treated with PBS (p=0.0110, n=6) but not CON-EV (p=0.2209, n=6, **Fig. 3A**). This pro-apoptotic effect is abrogated by 26% when CCA-EVs are permeabilized with 0.1% Triton X-100 before coculture with LLC cells (p=0.0024, n=6, **Fig. 3A**). Percent differences were calculated by dividing the means from each condition. Concomitantly, CCA-EVs significantly increased DNA fragmentation in LLC cells by 13% and 10% compared to cells treated with CON-EVs (p=0.0208, n=7) or PBS (p=0.0307, n=7, **Fig. 3B**) respectively. The induction of the pro-apoptotic effect was reduced by 18% when CCA-EVs were treated with Triton X-100 and proteinase K before co-culture with LLC cells (p=0.0307, n=7, **Fig. 3B**). To examine the effect of CCA-EVs on upstream mediators of cell death, we measured the expression of proteins associated with apoptosis such as p53, Bax, Bcl-2, cytochrome *c*, caspase-3, cleaved caspase-3 and AIF. CCA-EVs increased the expression of pro-apoptotic protein, Bax (p=0.0506, n=5, **Fig. 4A** and **4B**) by 25%, as well as Bax/Bcl-2 ratio (p=0.0349, n=5, **Fig. 4C**) by 60% in LLC cells compared to CON-EVs. The expression of other proteins remained unchanged (**Fig. 4A** and **4B**).

**Figure 3.**
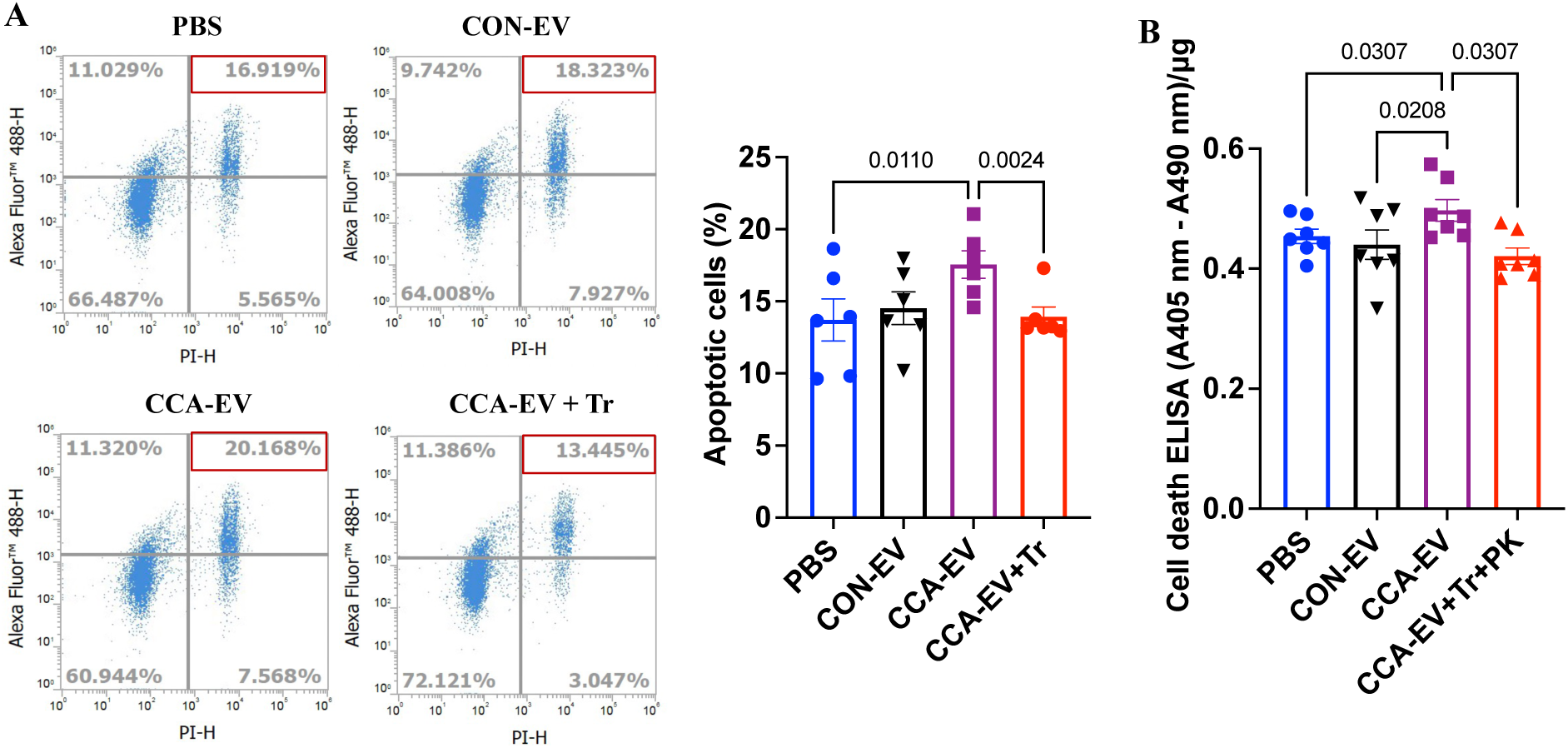
CCA-EVs increase markers of apoptosis in LLC cells. CCA-EVs were pretreated with 0.1% Triton X-100 with or without proteinase K (10 µg/mL, 1 h, 37 °C), and then co-cultured with LLC cells for 4 days. LLC cells were also treated with CON-EV and PBS for 4 days, after which the level of apoptosis was measured. **(A)** Representative images and quantification of Annexin V-FITC/PI staining assay. The percentage of apoptotic cells (double stained cells) are highlighted in red. **(B)** Cell death ELISA in treated LLC cells. Data were analyzed using a one-way ANOVA, with multiple comparisons corrected using Holm-Šídák or Dunn’s post hoc test, and expressed as scatter plots with mean (n=6-7). Exact p values for significant (p<0.05) results are shown. The p-values for non-significant data are not shown.

**Figure 4.**
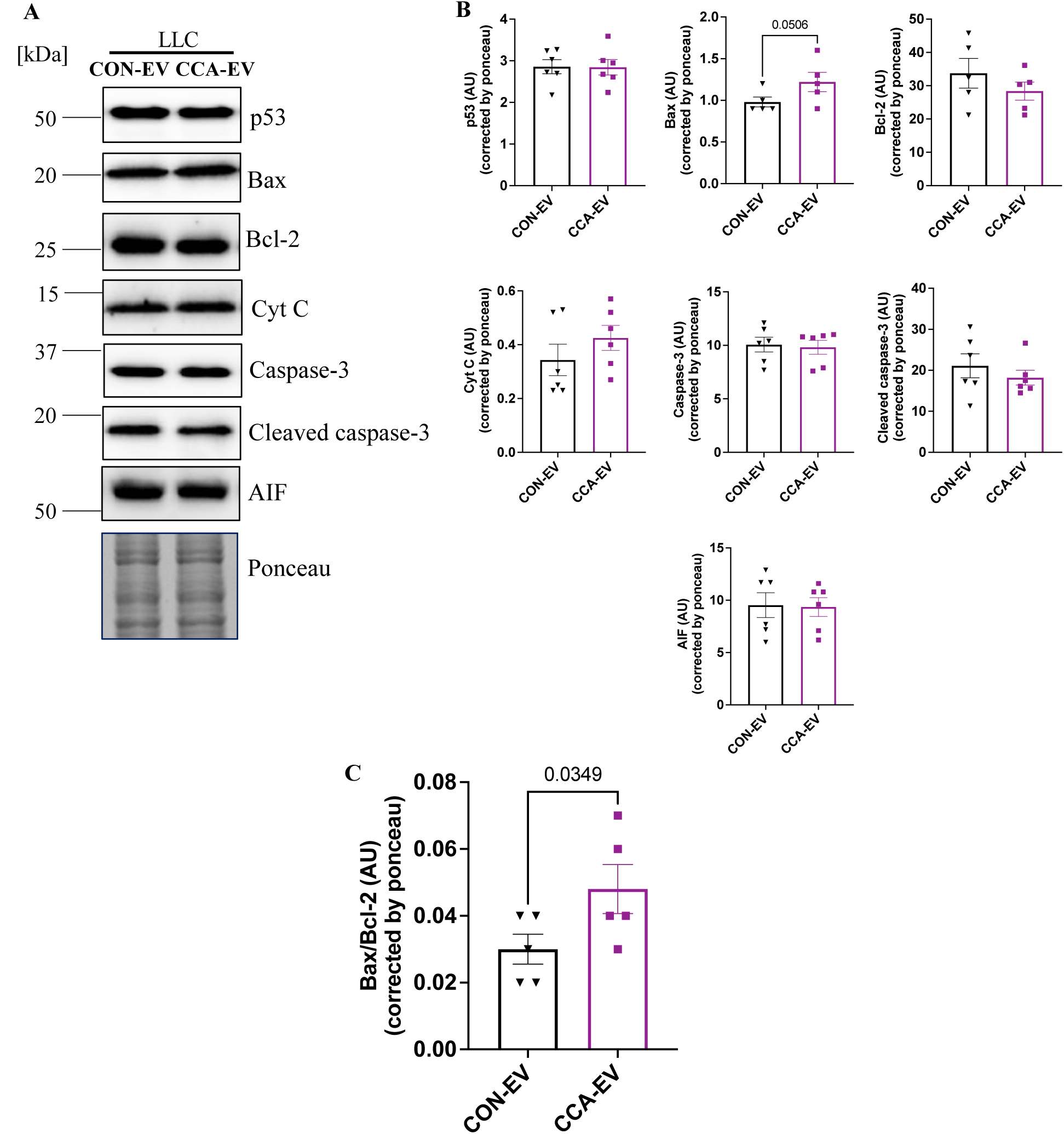
CCA-EVs increase the expression of some pro-apoptotic proteins in LLC cells. EVs were isolated and co-cultured with LLC cells after each bout of contractile activity as previously described. After the last day of treatment, cells were collected and assessed for protein expression **(A, B)** Representative blots and quantification of p53, Bax, Bcl-2, Cyt C, caspase-3, cleaved caspase-3, AIF, and **(C)** Bax/Bcl-2 ratio. Data were analyzed using an unpaired Student’s t-test and expressed as scatter plots with mean (n=5-6). Exact p values for significant (p<0.05) or close to statistically significant results is shown. The p-values for non-significant data are not shown.

### CCA-EVs induces senescence in non-small cell lung cancer cells

To ascertain whether CCA-EVs induced senescence in LLC cells in concert with apoptosis, we measured the level of senescence using SA-β-gal staining. CCA-EVs increased the percentage of SA-β-gal+ cells compared to cells treated with CON-EVs (p=0.0128, n=6) or PBS (p=0.0128, n=6), by 29% and 44%, respectively (**Fig. 5A**). Consistent with previous results, this pro-senescent effect was ameliorated by 56% when CCA-EVs were treated with Triton X-100 and proteinase K before coculture with LLC cells (p=0.0039, n=6, **Fig. 5A**). In addition, the expression of HMGB1 (p=0.0664, n=6) and p16 (p=0.0568, n=4), proteins involved in senescence, increased by 49% and 92% respectively in LLC cells treated with CCA-EVs compared to CON-EVs (**Fig. 5B**). The expression of Cyclin D1 remained unchanged (**Fig. 5B**).

**Figure 5.**
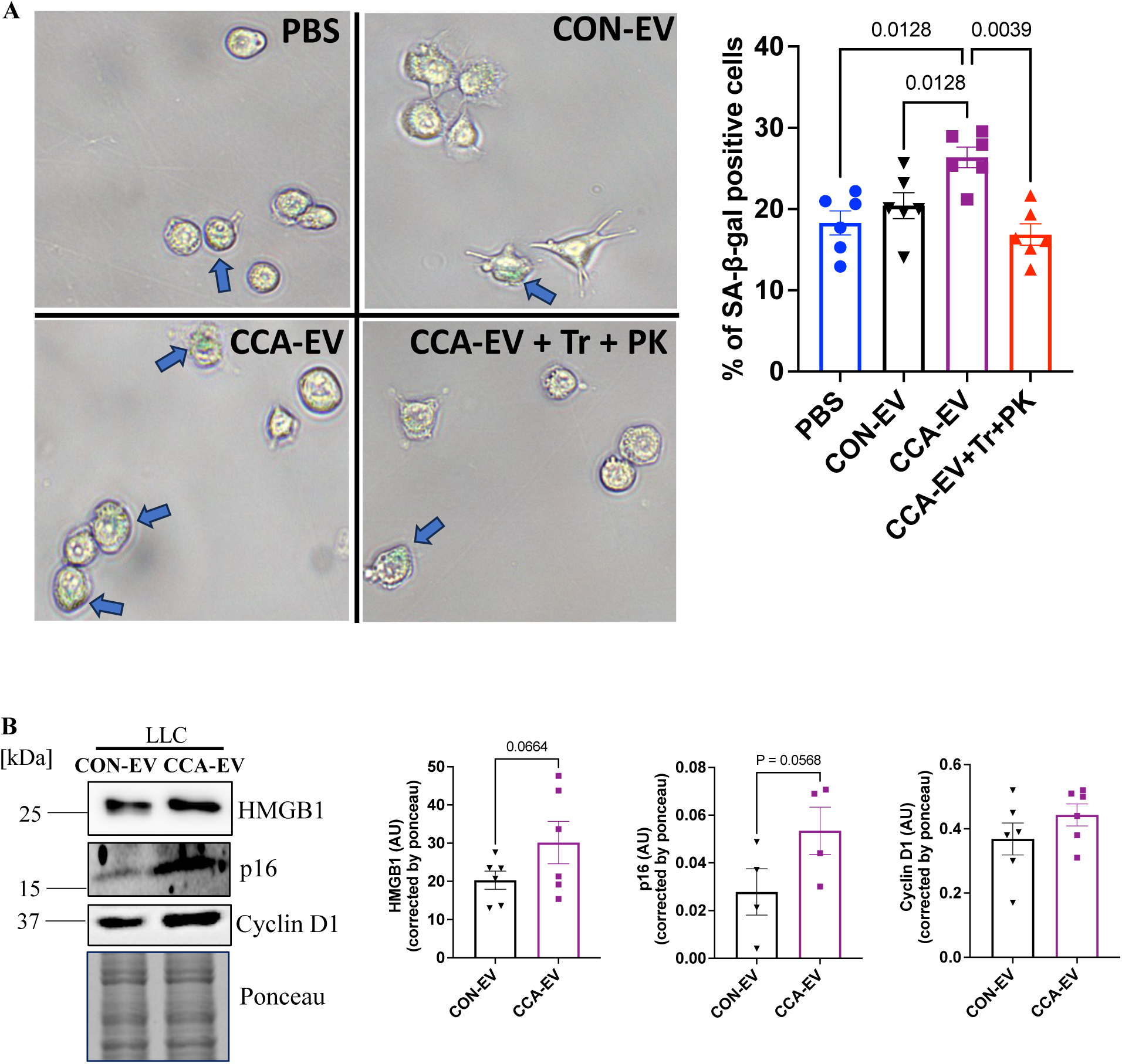
CCA-EVs induces senescence in LLC cells. CCA-EVs were pretreated with Triton X-100 with or without proteinase K, and then co-cultured with LLC cells for 4 days. LLC cells were also treated with CON-EV, CCA-EV and PBS for 4 days, after which the level of senescence was measured. **(A)** Representative images and quantification of SA-β-gal activity. Scale bar: 200 µm at 10X magnification, and **(B)** western blot analysis of senescence markers, HMGB1, p16, and cyclin D1 in treated LLC cells. Data were analyzed using a one-way ANOVA in panel A with multiple comparisons corrected using Holm- Šídák post hoc test, and by an unpaired Student’s t-test in panel B. Data are expressed as scatter plots with mean (n = 4-6). Exact p values for significant (p<0.05) or close to statistically significant results is shown. The p-values for non-significant data are not shown.

### CCA-EVs did not alter metastatic potential nor mitochondrial content in non-small cell lung cancer cells

In cancer, cell migration leads to invasion and metastasis, so we examined the effect of CCA-EVs on the migration of LLC cells using scratch assay. The results showed no change between cells treated CCA-EVs vs. CON-EVs or PBS. We also did not observe any difference when cells were treated with Triton + ProK-treated CCA-EVs (**Fig. 6A** and **6B**). Lastly, we examined the effect of CCA-EVs on mitochondrial content using MitoTracker Red, COX activity, and expression of complex I subunit (CI-NDUFB8), complex II subunit (CII-SDHB) and complex IV subunit (CIV-MTCO1) in LLC cells to delineate if EV treatment was modifying cellular metabolism. CCA-EV treatment did not change MitoTracker Red staining, a proxy for mitochondrial mass, (**Fig. 7A**), nor COX activity a well-standardized marker for mitochondrial biogenesis (**Fig. 7B**), or the expression of CI-NDUFB8, CII-SDHB, and CIV-MTCO1 (**Fig. 7C**) in LLC cells compared to CON-EVs.

**Figure 6.**
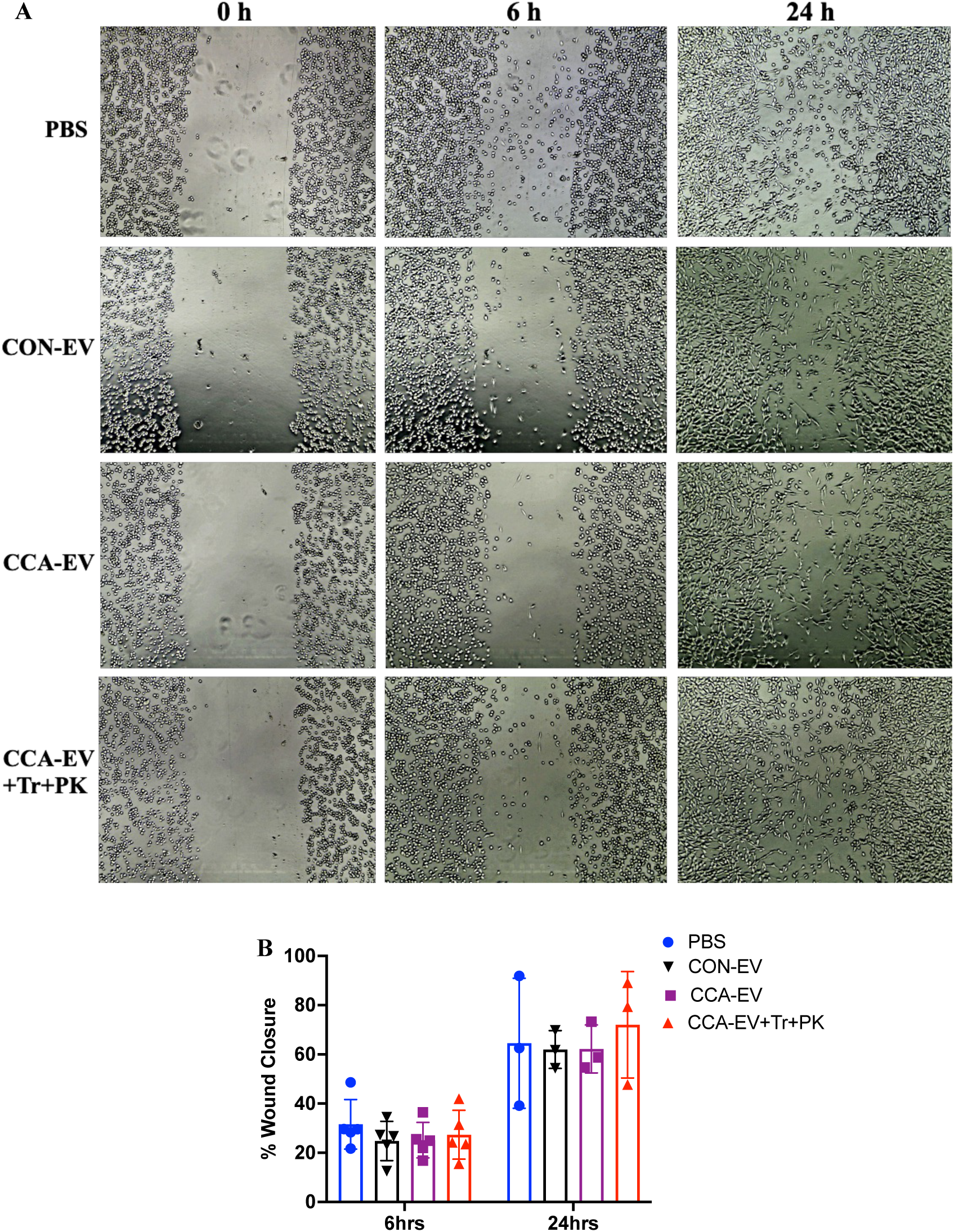
CCA-EVs did not affect migration in LLC cells. LLC cells were treated with CON-EV, CCA-EV and PBS for 4 days. CCA-EVs were also pretreated with Triton X-100 with proteinase K before co-culture with LLC cells for 4 days, after which the level of migration was measured using **(A, B)** Representative images and quantification of scratch assay in treated LLC cells at 0 h, 6 h and 24 h. 10X magnification. Data were analyzed using a two-way ANOVA, with multiple comparisons corrected using Bonferroni’s post hoc test (n = 3-5). The p-values for non-significant data are not shown.

**Figure 7.**
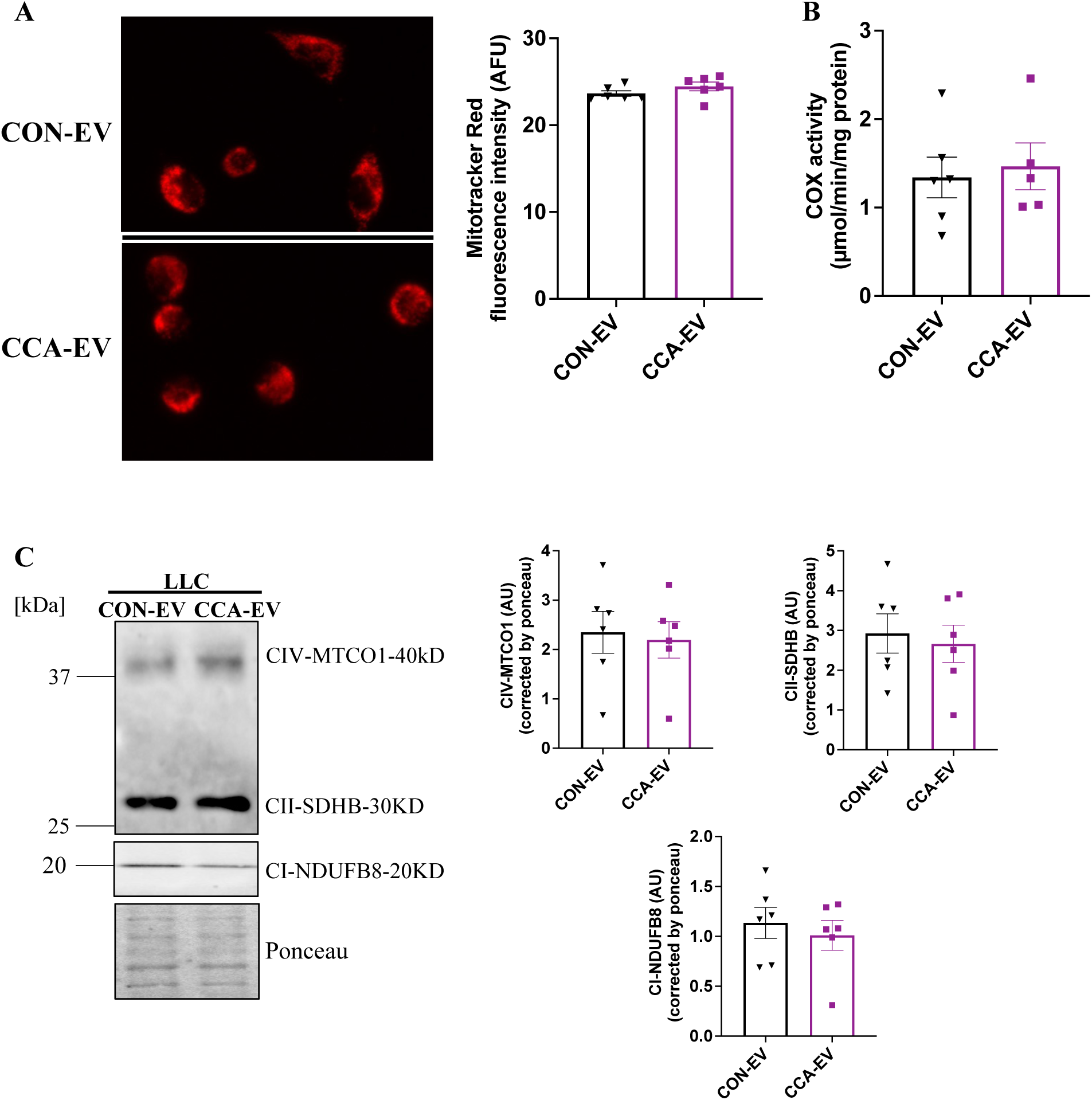
CCA-EVs did not change mitochondrial content in LLC cells. LLC were co-cultured EVs for 4 days. After the last day of treatment, markers for mitochondrial content were measured. **(A)** Representative fluorescent images and quantification of MitoTracker Red staining. Scale bar: 100 µm at 40X magnification. **(B)** Cytochrome *c* oxidase (COX) activity, and **(C)** western blot analysis of Complex I subunit (CI-NDUFB8), Complex II subunit (CII-SDHB) and Complex IV subunit (CIV-MTCO1) in LLC cells treated with CCA-EVs *vs*. CON-EVs. Data were analyzed using an unpaired Student’s t-test and expressed as scatter plots with mean (n = 5-6). The p-values for non-significant data are not shown.

### EV uptake by recipient LLC cells

Finally, to confirm uptake of EVs by recipient cells, we used Fluorogenic Membrane Probe, MemGlow 488, to label myotube-derived EVs as previously described [27]. MemGlow is a self-quenching dye that will not fluoresce until it integrates with a membrane, reducing the rate of non-specific binding. Labelled EVs were co-cultured with LLC cells for 24 h, after which we evaluated MemGlow+ EV uptake in fixed EV-treated LLC cells using confocal microscopy. We observed the presence of green fluorescence indicative of MemGlow+ EVs, co-localizing with the red fluorescence from rhodamine phalloidin (F-actin) showing EV uptake in recipient LLC cells (**Fig. 8**).

**Figure 8.**
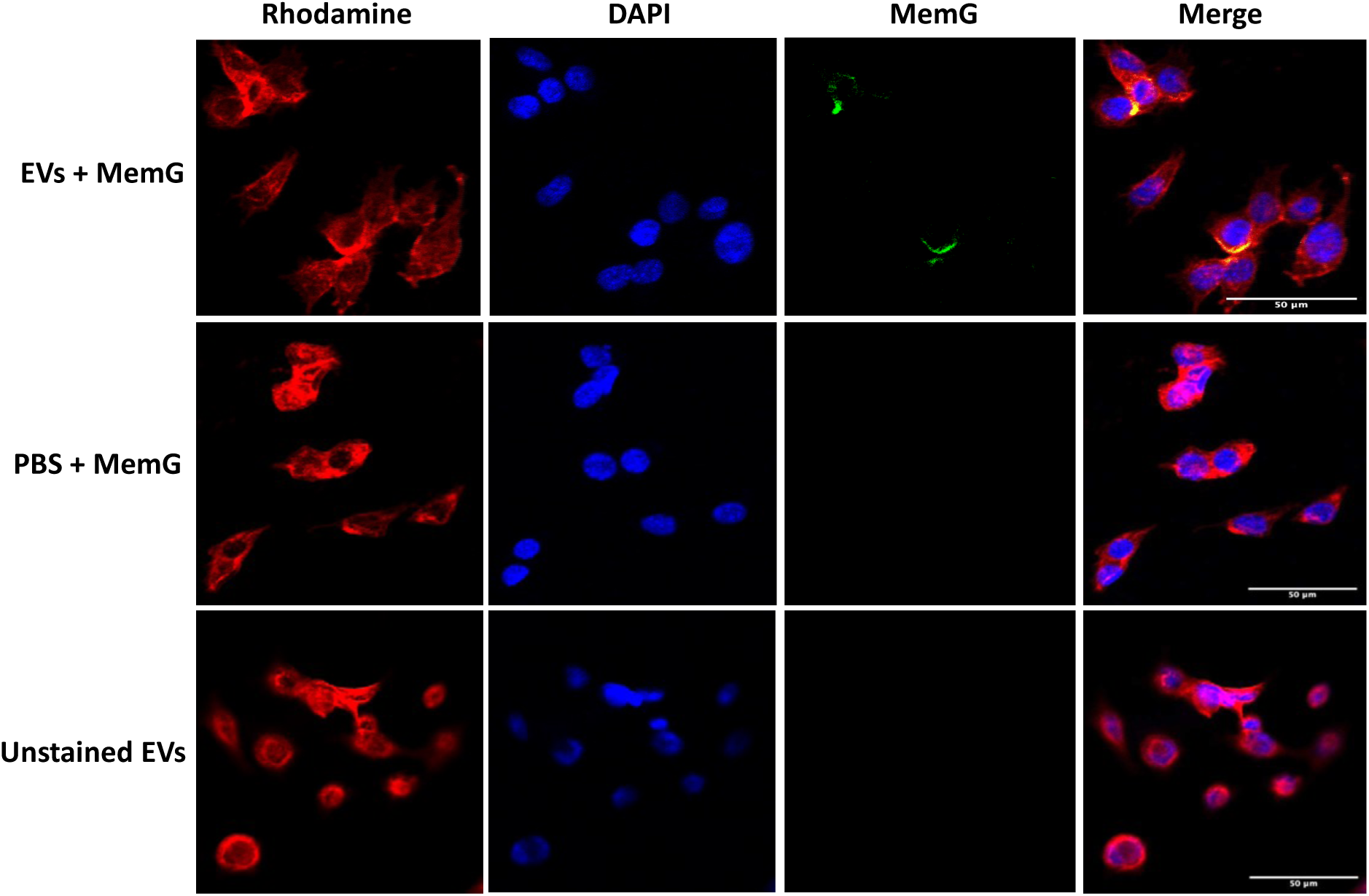
EV are taken up by recipient cells. MemG-labelled EVs (green) were used to treat LLC cells for 24 h, after which cells were fixed with 3.7% PFA, stained with Rhodamine phalloidin (red) and DAPI (blue), and imaged using confocal microscopy. PBS or unstained EVs were used as negative controls. Three representative images were taken per condition using the ZEN Pro software at 20X magnification, scale bar: 50 µm. Selected images demonstrate EV uptake by recipient cells.

## Discussion

Previous studies have shown that endurance exercise can suppress lung cancer growth in mice [35], [36]. A systematic review found exercise-conditioned human serum reduced cell viability in vitro in four types of cancers including lung cancer [37]. Another study illustrated that serum collected from humans after an acute bout of high intensity exercise led to the significant inhibition of survival and proliferation in non-small lung cancer cells [38]. Evidence indicates myokines including irisin, muscle-derived oncostatin and SPARC play a role in the exercise-mediated attenuation of cancer cell growth [11], [13], [14], but the exact mechanism(s) underlying the effects of endurance exercise on tumor biology is still poorly understood. Additionally, Sadovska et al. showed that EVs isolated from plasma of chronically exercised rats delayed the progression of prostate cancer [23]. However, the authors used plasma EVs, which originate different sources such as platelets, erythrocytes, endothelial cells, leukocytes, [20], [22], with little representation from skeletal muscle [25], [26]. The effect of skeletal muscle derived EVs after chronic activity on cancer cells remains unknown. Here, we illustrate for the first time that Skm-EVs released post-CCA can inhibit cell viability, increase cell death, and induce senescence in murine non-small lung cancer cell line, Lewis Lung Carcinoma (LLC) cells.

To evaluate the biological activity of Skm-EVs isolated post-chronic contractile activity, we treated LLC cells with CON-EVs or CCA-EVs isolated after each bout of contractile activity for 4 days. Using MemGlow 488, we confirmed that EVs can be taken up by recipient LLC cells. Next, we showed that CCA-EV co-culture treatment decreased cell count and viability using trypan blue exclusion but no significant changes were observed using the MTT assay. Both trypan blue and MTT assays are routine methods for evaluating cell viability [39], [40]. However, MTT reduction serves as an indicator of viable cell metabolism rather than a specific marker of cell proliferation [40], which could explain the disparate results. Indeed, Chung et al. showed a reduction in cell viability using Trypan blue assay but no significant changes using MTT assay [41]. In this study, although not significant, we observed an 11% decrease in cell viability with CCA-EV treatment using MTT. Since, MTT is an indicator of cell metabolism, it is likely that the 4-day EV treatment on LLC cells was too long to detect transient significant changes in the cell metabolic activity.

Mitochondrial-mediated apoptosis is a major apoptotic pathway that involves the activation or inhibition of several proteins including Bax (proapoptotic protein) and Bcl-2 (anti-apoptotic protein), mitochondrial membrane permeabilization, pro-apoptotic protein release (e.g. cyt c, AIF) and activation of the caspase cascade resulting in DNA fragmentation, the hallmark measurement of apoptosis [31], [42]. Tumor suppressor p53, an upstream regulator of the apoptotic process and a transcriptional activator of Bax [43], was not altered with CCA-EV treatment. CCA-EV treatment increased Bax expression but not Bcl-2, with a net increase in Bax/Bcl-2 ratio, a marker of apoptotic induction [44]. Cyt C, AIF, caspase-3, cleaved caspase 3 levels remained unchanged with CCA-EV treatment. However, we measured an increase in DNA fragmentation, the hallmark measurement of apoptosis. In concert, the number of apoptotic cells (Annexin V/PI positive cells), increased with CCA-EV treatment. Both of these are the end-stage markers of apoptosis. It is likely that we did not find differences in upstream signalling proteins as we measured the expression of these proteins in the whole cell lysates. Assessing the protein expression stratified by cellular localization, i.e., nuclear, cytoplasmic, vs. mitochondrial depots, will provide clearer insight into the upstream signalling pathway triggered by CCA-EVs. Additionally, we measured the effect on protein expression on day 5 of CCA-EV treatment. It is also plausible that alterations in protein expression of key pro-apoptotic proteins occurred earlier upon CCA-EV treatment.

Cellular senescence is a state of cell cycle arrest that serves as a protective mechanism to prevent proliferation in response to various stressors [45]. Senescence in cancer can be a double-edged sword, as it can be anti-or pro-tumorigenic under certain conditions [46]. Senescent cells in cancer can lead to the inhibition of cancer progression. Conversely, senescent cells can release senescence-associated secretory phenotype (SASP) factors, which can enhance tumor-promoting responses leading to cancer progression [46]. Here we showed that CCA-EV treatment increased the number of senescent or SA-β-gal-positive cells. HMGB1 protein expression increased in tandem with CCA-EV treatment, though not statistically significant. HMGB1 plays important roles based on its subcellular location. It acts as a DNA chaperone in the nucleus, exhibits cytokine-like function when released from the cell, and is to some extent expressed in the cytoplasm [47]. Additionally, cell death can cause the passive release of extracellular HMGB1. Evidence has also shown that HMGB1 is associated with senescence in cancer cells [48]. A previous study showed that cancer cells with a high HMGB1 expression selectively underwent senescence rather than apoptosis [49]. Higher HMGB1 protein expression with CCA-EV treatment supports the measured induction of senescence in this study. Similarly, CCA-EV treatment increased the expression of p16 and cyclin D1, which are upregulated during senescence [50], [51]. Overall, these results coupled with the cell viability and cellular apoptotic data indicate that CCA-EVs activated both cellular senescence and cell death, which likely contributed to the decrease in cell viability, as measured by trypan blue exclusion.

Migration or invasion of cancer cells leads to metastasis, thereby causing cancer progression [52]. Researchers have reported confounding results on the effect of exercise on cancer metastasis *in vivo*, the summary of evidence can be found here [10]. Briefly, evidence is mixed as some have reported an inhibition of metastasis, others an acceleration, while some have not found any effect on metastatic potential with exercise. Interestingly, exercise-conditioned serum from advanced-stage cancer patients inhibits the migration of pancreatic cancer cells [53]. Thus, we investigated the effect of CCA-EVs on the migratory potential of lung cancer cells. We did not observe any differences when with CCA-EV treatment compared to cells treated with CON-EVs. It is possible that a higher dose of CCA-EVs could provoke a significant effect on cell migration, and subsequently metastatic potential. A dose-response experiment with CCA-EVs on LLC is clearly warranted.

One of the hallmarks of cancer is deregulated cellular metabolism [54]. Evidence has shown that some cancer cells can reprogram their metabolism by using glycolysis as the main source of energy production (ATP) even in the presence of oxygen, a term known as aerobic glycolysis [55]. Even though cancer cells use the glycolytic pathway as their primary source of energy, they also utilize mitochondrial-mediated respiration for the production of ATP and metabolites, for lipid and nucleic acid metabolism, and in turn to permit rapid growth [56]. Indeed, a high mitochondrial content is positively associated with breast cancer progression and aggressiveness [57]. Thus, we investigated the effect of CCA-EVs on the mitochondrial content and function in LLC cells, but did not observe a difference when compared with cells treated with CON-EVs. Further investigation into mitochondrial oxygen consumption, and glycolytic capacity can inform if there is any modulation of cellular bioenergetics with CCA-EV treatment.

We pre-treated CCA-EVs with Triton X-100 with or without proteinase K before co-culture experiments with lung cancer cells, to determine if permeabilization with or without protein digestion would alter downstream effects of CCA-EV treatment. Interestingly, the effects seen in the lung cancer cells were diminished with the pre-treatment. This result is in line with our recent manuscript, where an increase in maximal respiration in myocytes treated with CCA-EVs was ameliorated when EVs were permeabilized and proteins digested [27]. This highlights that intact EVs are critical in delivering their cargo into recipient cells, and transmembrane, or membrane-associated proteins and/or corona of CCA-EVs execute the anti-tumorigenic effect of CCA. It is possible that the proteins on the surface of CCA-EVs are required for EV internalization through multiple pathways, and the removal of these proteins blocks the delivery of EV cargo to recipient cells. The proteins in the lumen of the EVs should not be ruled out altogether as it is possible that they are just not delivered internally into recipient cells, despite being present in permeabilized EV preparations. However, this explanation is speculative and further investigation is required identify putative membrane-associated and/or lumen cargo protein(s) that are differentially expressed with CCA and can perpetuate anti-cancerous signalling.

Our approach to mimic chronic exercise in myotubes using electrical pulse stimulation has been validated by our group. We showed an increase in mitochondrial biogenesis in myotubes that were chronically stimulated, 3 h/day for four consecutive days [27]. This approach is important helps to circumvent the challenge of using whole blood-based EV preparations to study the role of Skm-EVs. It is an efficient, fast method to induce the effect of exercise in cells as quickly as four days and interrogate the effect of contractile activity on Skm-EVs biological activity, without the heterogenous mixture of EVs in the plasma/serum. We also showed previously that CCA-EVs increased mitochondrial biogenesis but did not affect cell viability in C2C12 myoblasts [27]. Interestingly, in this current study, we treated lung cancer cells with the same CCA-EVs and observed opposite effects, a decrease in cell viability and no effect on mitochondrial biogenesis. This novel observation and begs the question as to why CCA-EVs have different biological activities in different cell types. Are different EV cargoes being delivered to different cell types? Is the effect mediated by recipient cell receptors? These are interesting questions that warrant further research. In summary, our data demonstrate for the first time that Skm-EVs derived post-CCA decrease cell viability, increase cell death, and induce senescence in NSCLC cells, and this effect is likely mediated by the membrane-bound protein cargo of EVs. Identifying the specific EV protein(s) involved in this process will help understand the mechanisms of the anti-tumorigenic effects of exercise.

## Author Contributions

P.O.O. and T.F.G.S., performed experiments in the current study, analyzed data, and created figures. P.O.O. and A.S. helped write and revise the manuscript. K.J.M, A.R.W., and J.W.G. provided technical and theoretical expertise to complete the work. All authors were involved in manuscript revisions. A.S. designed the project, and helped synthesize data, create figures, write, and edit the manuscript. A.S. is the corresponding author and directly supervised the project. All authors have read, edited and agreed to the published version of the manuscript.

## Acknowledgement

The authors would like to thank Benjamin Bydak for providing assistance with using the IonOptix C-PACE EM; Samira Seif for providing technical support; Jared Field for helping with the Annexin V/PI staining using flow cytometer; Muhammad Talha Mustafa for assisting with the immunoblotting of pro-apoptotic proteins; Martha Hinton for helping with the EV uptake experiments using confocal microscopy; and Emily Turner-Brannen for providing assistance with the COX activity experiment using Agilent BioTek Cytation.

